# Multiple trade-offs between defense and competitiveness traits in a planktonic predator-prey system

**DOI:** 10.1101/2022.05.02.490268

**Authors:** Tom Réveillon, Lutz Becks

## Abstract

Predator-prey interactions play a central role in community dynamics and depends on the covariation of traits of the interacting organisms. Intraspecific trade-off relationships between defense and competitiveness traits are important for understanding trophic interactions. However, quantifying the relevant traits forming defense-competitiveness trade-offs and how these traits determine prey and predator fitness remain major challenges. Here, we conducted feeding and growth experiments to assess multiple traits related to defense and competitiveness in 6 different strains of the green alga *Chlamydomonas reinhardtii* exposed to predation by the rotifer *Brachionus calyciflorus.* We found large differences in defense and competitiveness traits among prey strains and negative relationships between defense and competitiveness traits. Because we compared trait differences among strains whose ancestors previously evolved in controlled environments where selection favored defense or competitiveness, these negative correlations suggest the presence of a trade-off between defense and competitiveness. This trade-off was found for multiple combinations of defense and competitiveness traits. Furthermore, the differences in traits translated into differences in prey and predator fitness, which demonstrated the contribution of intraspecific trade-offs for predicting the outcome of predator-prey interactions.

## Introduction

Predator-prey interactions and their consequences on population dynamics are primarily determined by combinations of traits driving the performance of organisms (Colina et al. 2016). Understanding the mechanisms linking traits of predators and prey to the strength of trophic interactions is therefore crucial for assessing the functioning of food webs (Rossberg et al. 2010; Gravel et al. 2016; Laigle et al. 2018; Litchman et al. 2021). Traits involved in defense against predation (i.e., *sensu* protection) and competitiveness (i.e., *sensu* reproduction) are intimately related to the fitness of organisms and mediate trophic interactions (Colina et al. 2016). Prey can exhibit a large range of defense mechanisms against predators, which interfere with different steps of the predation sequence (Weiss et al. 2012; Bateman et al. 2014). These traits affect predator the predator by reducing prey detection and consumption (Rall et al. 2012) via altered attack rate (i.e., the rate of prey encounter), handling time (i.e., the time spent on prey manipulation and ingestion) or gape-size limitation and thus have consequences on the fitness of predators (Long and Hay 2006).

Organisms cannot optimize all functions contributing to fitness simultaneously due to genetic and/or energetic constraints and trade-off relationships between traits can emerge (Stearns 1989). Trade-offs between defense and competitiveness traits are expected (Yoshida et al. 2003; 2004; Becks et al. 2010; Sunda and Hardison 2010; Kasada et al. 2014; Ehrlich et al. 2017; Pančić and Kiørboe 2018; Cadier et al. 2019) and may explain the diversity of fitness strategies (Agrawal 1998; Strauss et al. 2002). Competitiveness costs can arise from the energy investment in expressing anti-grazing defenses due to a reduction in resource acquisition and utilization (Halsey and Jones 2015), and thus in reproduction (Smith et al. 2014). Therefore, trade-off relationships between defense and competitiveness traits can affect trophic interactions by influencing both prey and predator population growth (Wood et al. 2020). However, such trade-offs are complex to assess because multiple traits can contribute to these relationships. Evaluating suitable traits describing defense-competitiveness trade-offs using a multiple traits approach can give important insights on the presence and the role of these trade-offs for population dynamics and energy transfers in food webs.

Plankton communities are characterized by strong trophic interactions and trade-offs between defense and competitiveness traits play an important role in regulating these interactions (Litchman et al. 2007; 2015; Ehrlich et al. 2020). However, there are few empirical data on defense-competitiveness trade-offs and the existing data should be interpreted with caution. First, many studies assume fixed traits and trade-offs at the genus or species level (Bruggeman 2011). However, this assumption has been shown to be often invalid (Züst and Agrawal 2017) as intraspecific traits and trade-offs can be ecologically and evolutionary important (Yoshida et al. 2004; Becks et al. 2010). Second, defensive and competitive abilities of phytoplankton can be assessed by multiple traits (Fleischer et al. 2018) and the relevant traits constituting a trade-off are often unknown. Previous studies used different traits as proxies for defense and competitiveness and thus different trait correlations. This makes comparisons of trade-offs across studies difficult and raises the question of whether different traits can be used for studying the role of trade-offs for planktonic communities. Third, correlations between traits might not reflect the absence or the presence of a causal trade-off relationship due to confounding factors when traits are independently affected by the same environment (Bohannan et al. 2002; Fry 2003). False-positive (i.e., negative correlation) or false-negative (i.e., no or positive correlation) trade-offs could for example arise from variation in a third correlated trait which was unaccounted (Edwards et al. 2011; 2013). Finally, it remains unclear whether a negative correlation represents a trade-off or simply a signature of differential selection without knowing whether evolution could optimize both traits simultaneously (Fry 2003; Fuller et al. 2005). Perhaps one of the most thorough demonstrations of trade-offs is when a trait under selection is optimized at the expense of a trait not under selection and vice versa (Kassen 2002).

Here, we investigate the intraspecific trade-off relationship between anti-grazing defense and competitiveness traits for 6 different prey genotypes (hereafter: strains) of the green alga *Chlamydomonas reinhardtii* exposed to the rotifer predator *Brachionus calyciflorus.* Algal strains were isolated from a previous experimental evolution study where they were grown under selection for either defense or competitiveness (Bernardes et al. 2021). They differed in their morphologies (i.e., single or colonial cells) as well as defense (i.e., measured as reduction in predator growth rate) and growth (i.e., measured as the prey growth rate). We quantified prey defense against a predator by measuring functional responses (i.e., the ingestion rate as a function of prey density) on prey strains to derive the attack rates and the handling times using mechanistic models, and also by measuring a range of size and shape characteristics and the carbon to nitrogen ratio of prey strains to link prey morphology and stoichiometry to the trophic interaction with the predator. We further quantified prey competitiveness by measuring prey growth in limiting nitrate conditions to derive the maximum growth rate, the nitrate affinity and the nitrate half-saturation constants using mechanistic models. We tested for the trade-off relationship between defense and competitive traits, whether it was present for all trait combinations, and how the trait and trade-off differences affected population growth (i.e., fitness) of the algal prey strains and the rotifer predator.

## Material and methods

We used 6 strains of the green alga prey *Chlamydomonas reinhardtii* (Dang 1888) with known recent evolutionary history which were derived from isolates obtained from the Chlamydomonas Resource Center (University of Minnesota, USA). Isolates originated from an experimental evolutionary study in which different monoclonal naive ancestors were selected either for defense (exposed to the rotifer *Brachionus calyciflorus*) or competitiveness (not exposed) for 6 months (∼500 generations) in semi-continuous cultures (Tab. S1; Bernardes et al. 2021). For the present study, a single colony from each evolved strain was sampled from solid media and transferred to tissue culture flasks filled with 50 mL of sterile growth medium with 800 µmol NO_3_^-^ L^-1^. Flasks were exposed to continuous light (200 µmol photon m^-2^ sec^-1^) and shaking (120 rpm) at 20°C for 2 weeks before the assays, which allowed the cultures to reach relatively high densities. We used a clonal line of the rotifer predator *Brachionus calyciflorus* Pallas (Pallas 1766) derived from an isolate sampled in Milwaukee harbor (Bennett and Boraas 1989). This clonal line lost the ability to reproduce sexually and the possibility for evolutionary changes through genetic mixing (Fussmann et al. 2003). Rotifer stocks were kept in 1 L bottles filled with sterile growth medium with 800 µmol NO_3_^-^ L^-1^ (Barreiro and Hairston 2013) and fed with the nutrient-rich chlorophyte alga *Monoraphidum minutum* obtained from the Culture Collection of Algae (University of Göttingen, Germany).

### Prey morphology

To assess the variation in morphological traits among strains, *C. reinhardtii* strains were sampled (100 µL) from the culture flasks and transferred into 1.5 mL tubes. Samples were mixed and imaged using image flow cytometry (Amnis^®^ ImageStreamX Mk II). A collection of 5000 images each showing a single cell or a cell colony was acquired for each strain with a 20X magnification using the Cy5 channel (642 nm) and the autofluorescence signal of algal cells. We chose 9 morphological features (Tab. S2) that were calculated per image using the instrument analysis software (Amnis IDEAS^®^, see *Supplementary Materials Appendix 2*). Features were divided in 2 categories: cell size and cell shape. We used a principal components analysis (PCA) on the morphological features estimated on individual cell images to identify the most relevant morphological traits for characterizing strains. The PCA was run on the image features and the 2 first dimensions explaining most of the variance in the data were selected (D_1_ = 69.43%, D_2_ = 18.67%). We performed parametric correlations between 2 relevant features associated to cell size (feature: *area*) and cell shape (feature: *roundness*) according to contributions to the PCA dimensions. Statistical differences among strains for these 2 features were tested using Kruskal-Wallis and Dunn post-hoc tests. The position of features on the PCA dimensions (Fig. S4) and the correlations (Fig. S5) showed a significant relationship between *area* and *roundness* of prey strains. We chose to include the feature *area* representing prey particle size, which includes single cells or cell colonies, as a proxy trait for anti-grazing defense (Lürling 2021).

### Prey stoichiometry

To assess the variation in stoichiometric traits among strains, *C. reinhardtii* strains were sampled (10 mL) from culture flasks and centrifuged at 2000 rpm for 10 min at 20 °C. The culture medium was removed and the pellets were suspended in 5 mL of fresh culture medium at a density of 5×10^5^ cells mL^-1^. Carbon and nitrogen cell masses were measured following a combustion oxidation method (Bremner 1965). Algae were filtrated on 0.2 µm-mesh filters that were pre-calcinated for 2 h in a muffle furnace at 550°C to remove carbon. Filters were dried for 24 h in an oven at 60°C and stored for 24 h in a desiccator dome at 20°C. Dried filters were then packed in tin capsules and oxidated by combustion at 1020°C into a gas chromatographer to estimate carbon and nitrogen masses (Hekatech^®^ Euro Elemental Analyzer). The experimental designs included 3 replicates for each strain and 3 control treatments of culture medium. Cell mass of each element (µg cell^-1^) was calculated as: M = (M_S_ – M_C_) / (D × V), where *M_S_* is the total element concentration in the algae solution (µg mL^-1^), *M_C_* is the total element concentration in the control solution (µg mL^-1^), *D* is the cell density (cells mL^-1^) and *V* is the volume of the filtrate (mL).

### Prey defense

Prior to the experiment, *C. reinhardtii* strains were sampled from culture flasks and centrifuged at 2000 rpm for 10 min at 20 °C. The culture medium was removed and the pellets were suspended in nitrate-free medium at a density of 1×10^6^ cells mL^-1^ for 24 h to significantly slow down algal growth during the exposure to predation. At the start of the experiment, strain solutions were diluted to 10 different cell densities (1×10^4^ to 1×10^6^ cells mL^-1^). These dilutions were made in 2 mL tubes by mixing the strains with nitrate-free medium and 200 µL were transferred into wells of 96 well plates. The experimental design included 3 replicates per cell density for each strain for a control treatment without exposure to predation to estimate the initial densities in the wells and for a predation treatment to estimate the final densities after exposure to predation. For the predation treatment, 4 adult rotifers without carrying eggs were introduced per well and allowed to feed for 8 h at 20°C under continuous light. The duration of the experiment did not allow for rotifer reproduction to maintain a constant predation pressure. Afterwards, cells were fixed by adding 10 µL of Lugol solution per well and plates were stored at 4°C in the dark to allow cells to settle to the bottom of the wells. Initial and final cell densities were assessed by taking 21 images per well using a Cy5 filter set and the autofluorescence of the algal cells (642 nm) under a 10x magnification using a high content microscope (ImageXpress^®^ Micro 4 High-Content Imaging System). Images were analyzed and cell densities were calculated using a custom module within an analysis software (MetaXpress^®^ High Content Image Acquisition and Analysis, *Supplementary Materials Appendix 1*). Ingestion rates were calculated as i_R_ = (C_RI_ – C_RF_) / (t × B_C_), where *C_RI_* is the mean initial cell density of replicate wells from the control treatment (cells mL^-1^), *C_RF_* is the final cell density of each well from the predation treatment (cells mL^-1^), *t* is the time of feeding (sec) and *B_C_* is the number of rotifers per well (individuals).

To obtain predator funtionnal responses and derive predator attack rates (*a_i_*) and handling times (*h_i_*), the mean ingestion rate of the 3 replicates was calculated over the range of cell densities for each strain. Following the disc equation (Holling 1959), functional responses were expressed as mean ingestion rate of the predator (*i_Ri_*) as a function of prey densities (*C_Ri_*) and predator ingestion parameters. We fitted the Holling type I, II and III (Holling 1965) and the Ivlev type II models (Ivlev 1961) for each strain using non-linear least squares regressions and compared their AIC values (Tab. S2). The best fitting model was the Holling type II model for strains C_R2_, C_R3_, C_R6_, C_R7_ (Eq. 1) and Holling type III model for strains C_R1_ and C_R4_ (Eq. 2):

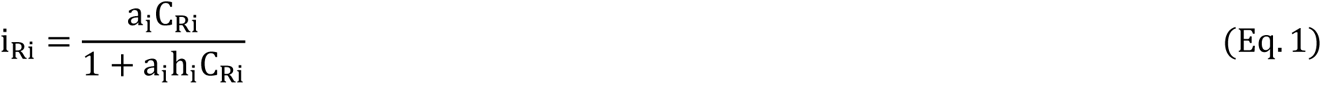

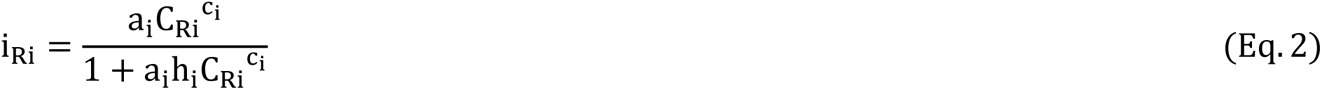

where *a_i_* is the attack rate of the predator on strain *i* (10^-6^ mL sec^-1^), *h_i_* is the handling time of the predator on strain *i* (sec), *c_i_* is the prey density exponent of the predator on strain *i* and *C_Ri_* is the cell density of strain *i* (cells mL^-1^). Because prey density declined over the duration of the experiment due to predation, we incorporated the decrease in prey density for the calculation of ingestion rates (Rosenbaum and Rall 2018). For this, we computed of system of differential equations to simulate dynamics of prey density over time and solved the Holling type II or type III model for each strain using an iterative maximum likelihood estimation (hereafter: prey density-corrected ingestion rate). We then derived the attack rates *a_i_* and handling times *h_i_* of each strain from the fitted models (see *Supplementary Materials Appendix 3*).

### Prey competitiveness

Following the same protocol described above, *C. reinhardtii* strains were concentrated 24 h prior to the experiment. We measured growth under 7 different nitrate concentrations: 6 low concentrations (1−10 µmol NO_3_^-^ L^-1^) and 1 high concentration (100 µmol NO_3_^-^ L^-1^) to calculate growth rates, nitrate affinity constants and nitrate half-saturation constants. At the start of the experiment, strain solutions were diluted to an initial cell density of 1×10^3^ cells mL^-1^. Growth assays were made in 96 well plates in a volume of 200 µL by mixing the strains (20 µL) and medium (180 µL). We used a disruptive sampling method: a well plate was assigned to each day and cells were fixed daily in one of the replicated well plates by adding 10 µL of Lugol solution per well. The experimental design included 3 technical replicates per nitrate concentration for each strain. Plates with fixed cells were stored at 4°C in the dark for 24 h to allow cells to settle to the bottom of the wells. Cell densities were assessed as described above and growth curves were estimated over 8 days.

To derive prey maximum growth rates (*r_Cimax_*), nitrate affinity constants (*f_Ci_*) and half-saturation constants (*K_Ci_*), we computed non-linear least squares regressions on prey growth rates as a function of nitrate concentrations for each strain using the Monod model (Eq. 4):

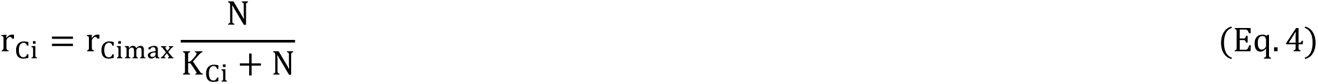

where *r_Cimax_* is the maximum growth rate of strain *i* among nitrate concentrations (day^-1^), *K_Ci_* is the nitrate half-saturation constant of strain *i* (µmol NO_3_^-^ L^-1^) and *N* is nitrate concentration (µmol NO_3_^-^ L^-1^). Nitrate affinity constants of strains were calculated as f*_C_*_i_ = r_Cimax_ / K_Ci_. To obtain prey growth rates (*r_Ci_*), we computed linear regressions on the growth curves for each strain and nitrate concentration. We applied these regressions on the first 3 days of the growth curves where nitrate concentrations were expected to be non-limiting for all strains.

To obtain prey growth curves at different nitrate concentrations, the mean cell density of the 3 replicates was calculated for each strain, nitrate concentration and day. Following previous studies (Paine et al. 2012; Malerba et al. 2018), we chose 2 non-asymptomatic growth models which assume an unlimited cell growth over time (i.e., linear and exponential models) and 3 asymptotic growth models which assume a limited cell growth with a maximum cell density reached over time (i.e., logistic, gompertz and mortality models). We fitted the growth models to each combination of strain and nitrate concentration using non-linear least squares regressions on cell densities as a function of time and compared their AIC values (Tab. S4). The best fitting model was the logistic model for strains C_R1_, C_R2_, C_R3_, C_R6_ and C_R7_ (Eq. 3) and the mortality model for strain C_R4_ (Eq. 5):

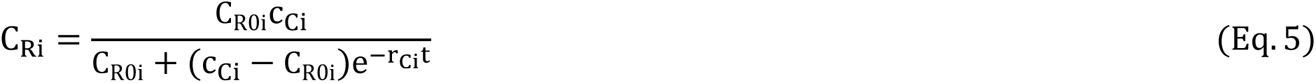

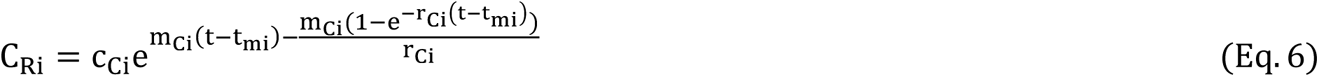

where *C_Ri_* is the cell density of strain *i* at time *t* (cells mL^-1^), *C_R0i_* is the initial cell density of strain *i* (cells mL^-1^), *c_Ci_* is the asymptotic cell density of strain *i* (cells mL^-1^), *r_Ci_* is the intrinsic growth rate of strain *i* (cells day^-1^), *m_Ci_* is the decreasing slope following maximum cell density of strain *i* (cell day^-1^), *t* is the time (day) and *t_mi_* is the time of maximum intrinsic growth rate of strain *i* (day).

### Prey fitness

We used the prey growth curves and the prey growth rates under the highest nitrate conditions in the experiment described above (100 µmol NO_3_ ^-^ L^-1^) to infer the differences in prey fitness among *C. reinhardtii* strains.

### Predator fitness

Prior to the experiment, *C. reinhardtii* strains were sampled from culture flasks and concentrated in nitrate-free medium at a density of 1×10^6^ cells mL^-1^ as described above. At the start of the experiment, strain densities were estimated and diluted to 5×10^5^ cells mL^-1^ in 1 mL of a nitrate-free medium in 24 well plates. The nitrate-free medium prevented prey growth and the high initial cell density provided sufficient resource for maximal predator growth. We introduced 2 juveniles and 3 adults *B. calyciflorus* carrying 1 or 2 eggs per well to maintain a homogeneous age structure in the initial population. The experimental design included 5 replicates for each strain. Rotifer individuals were daily counted using a stereomicroscope and growth curves were estimated over 5 days.

To obtain predator growth curves, the mean rotifer density of the 5 replicates was calculated for each strain and day. We chose 2 non-asymptotic growth models (i.e., linear and exponential models) and 2 asymptotic growth models (i.e., logistic model and mortality models). We fitted the growth models using non-linear least squares regressions on rotifer densities as a function of time and compared their AIC values (Tab. S5). The best fitting model was the mortality model for strains C_R1_, C_R2_, C_R3_ and C_R4_ (Eq. 4) and the logistic model for strains C_R6_ and C_R7_ (Eq. 3). We computed linear regressions on the growth curves for each strain to obtain the predator growth rates (*r_Bi_*). We used the predator growth curves and the predator growth rates to infer the differences in predator fitness among *C. reinhardtii* strains.

### Statistical analysis

All statistical analyses were computed using the R software v.4.0.4 (R Development Core Team 2020). Linear regressions and non-linear least squares regressions were computed using the *lm* and *nls* functions from the ‘lme4’ R package. Confidence intervals for non-linear least squares regressions were computed using the *predictnls* function from the ‘propagate’ R package. Models were compared using the *AIC* function from the ‘stats’ R package. Maximum likelihood regressions were computed using the *mle2* function from the ‘bbmle’ R package. Parametric correlations were computed using the function *cor* from the ‘stats’ R package. The principal components analysis was computed using the *PCA* function from the ‘FactoMineR’ R package. Graphics were obtained using the *ggplot* function with associated subfunctions from the ‘ggplot2’ R package.

## Results

### Prey traits

The defense of *C. reinhardtii* strains was defined based on the maximum ingestion rate of the predator *B. calyciflorus* and ranged from defended (low ingestion) to undefended (high ingestion). *B. calyciflorus* ingestion rates differed among strains (Fig. 1a). Strains C_R1_, C_R2_ and C_R3_ were classified as more defended due to the low maximum ingestion rates exhibited by the predator while strains C_R4_, C_R6_ and C_R7_ were classified as less defended due to the high maximum ingestion rates exhibited by the predator. The comparison of functional response models revealed that the ingestion rates of *B. calyciflorus* on *C. reinhardtii* strains were better described by non-linear saturating models (Tab. S3 and Fig. S1a). The differences in defense between strains were evident by the shapes of the functional responses (Fig. 1a) and lower attack rates but higher handling times of *B. calyciflorus* on more defended strains compared to less defended strains (Tab. 1). The competitiveness of *C. reinhardtii* strains was classified based on the maximum growth rate and ranged from less competitive (low growth) to more competitive (high growth). Growth parameters related to nitrate differed among *C. reinhardtii* strains (Fig. 1b). More defended strains C_R1_, C_R2_ and C_R3_ were classified as less competitive due to the low nitrate affinity and the high nitrate half-saturation constants while less defended strains C_R4_, C_R6_ and C_R7_ were classified as more competitive due to the high nitrate affinity and the low half-saturation constants (Tab. 1).

**Figure 1:**
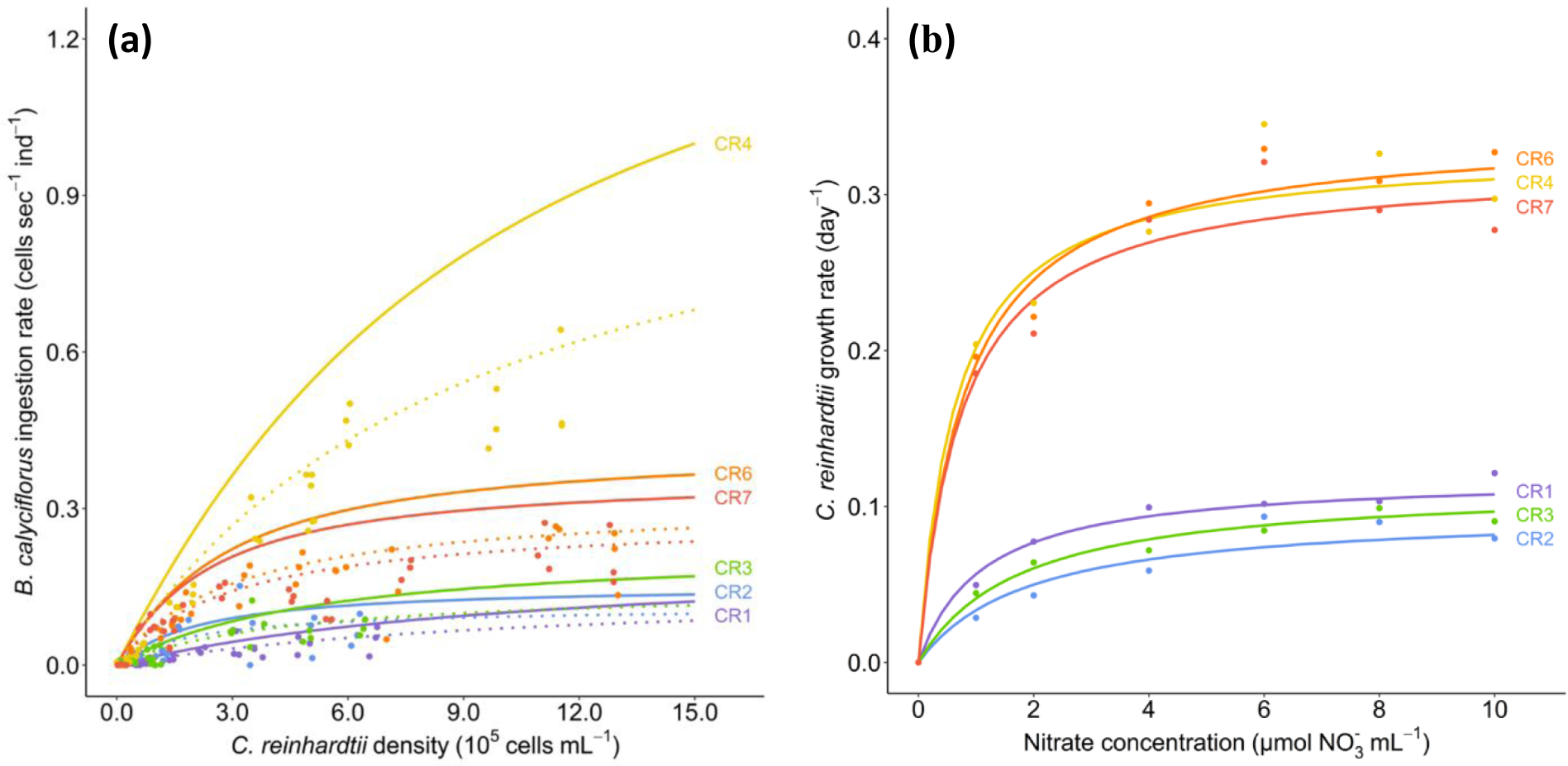
Defense and competitiveness of *C. reinhardtii* strains. (a) Functional responses of *B. calyciflorus* showing ingestion rate as a function of cell density of *C. reinhardtii* strains. Predicted functional response curves (lines) were estimated from the 3 replicates for ingestion rates (dots) for raw data (dotted lines) and prey density-corrected data (solid lines). (b) Monod curves of *C. reinhardtii* strains showing growth rate as a function of nitrate concentration (1−10 µmol NO_3_^-^ L^-1^). Predicted curves were estimated from the growth rate at each nitrate concentration (dots). Predicted curves with lower and upper boundaries using standard errors are shown in the *Supplementary Materials* (Fig. S1a and S1b).

**Table 1:**
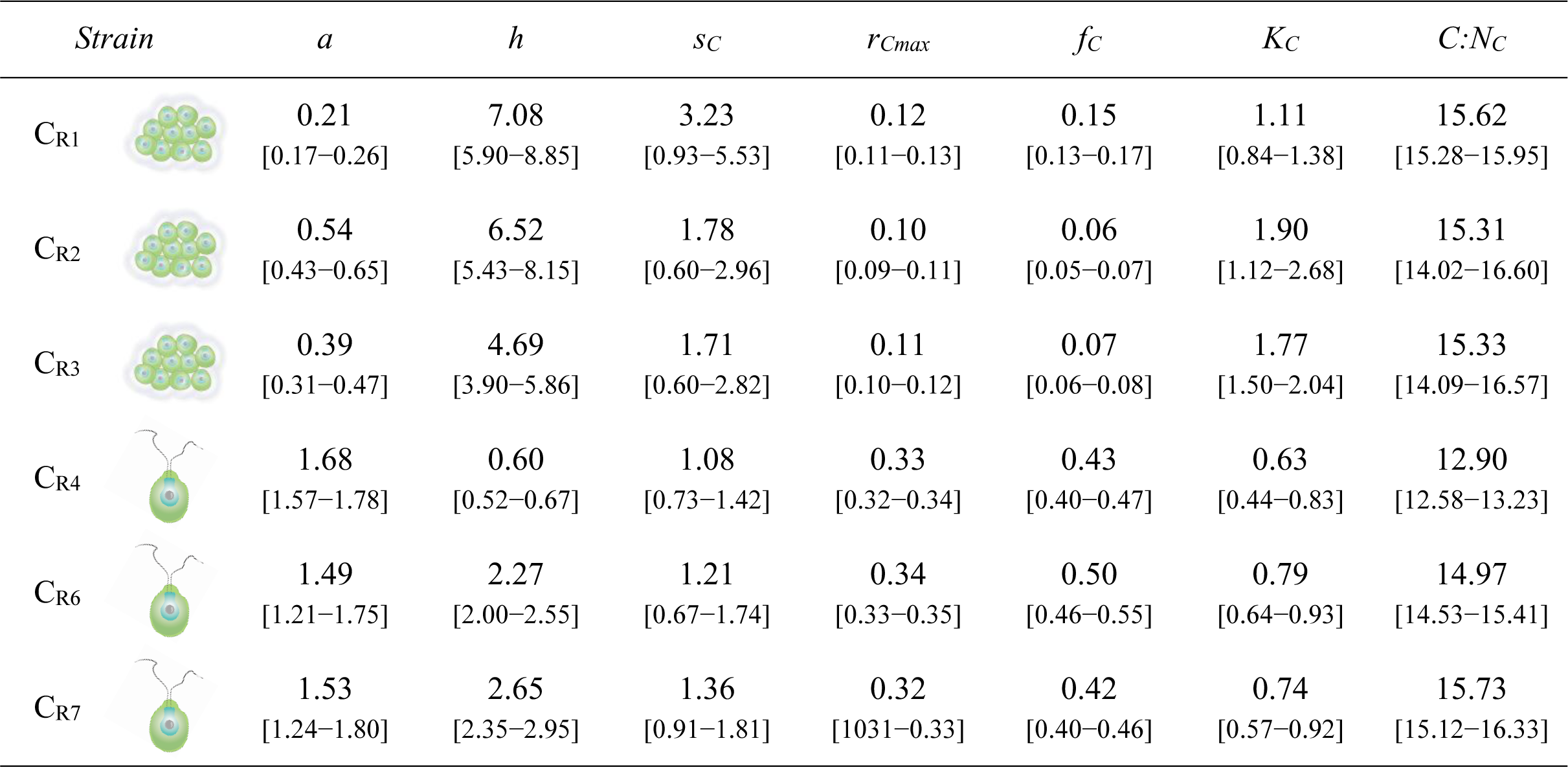
Estimates of defense and competitiveness traits [± standard deviation] for the 6 different *C. reinhardtii* strains. Traits were combined pairwise to construct the different defense-competitiveness trait spaces (Fig. 2). Anti-grazing defenses were represented by *B. calyciflorus* attack rate (*a* in 10^-6^ mL sec^-1^), handling time (*h* in sec), *C. reinhardtii* particle size (*s_C_* in 10^3^ µm^2^) and carbon to nitrogen ratio (*C:N_C_*). Competitiveness ability was represented by *C. reinhardtii* maximum growth rate (*r_Cmax_* in day^-1^), nitrate affinity constant (*f_C_* in µmol NO_3_^-^ L^-1^ day^-1^) and nitrate half-saturation constant (*K_C_* in µmol NO_3_^-^ L^-1^). Strains exhibited different morphologies in terms of cell clumping: clumping strains were C_R1_ (large clumps [10−30 cells]), C_R2_ (medium clumps [5−10 cells]) and C_R3_ (medium clumps [5−10 cells]) while non-clumping strains were C_R4_ (small single cells), C_R6_ (small single cells) and C_R7_ (small single cells).

| Strain | *a* | *h* | *s<sub>C</sub>* | *r<sub>Cmax</sub>* | *f<sub>C</sub>* | *K<sub>C</sub>* | *C:N<sub>C</sub>* |
| --- | --- | --- | --- | --- | --- | --- | --- |
| C<sub>R1</sub> | 0.21<br>[0.17–0.26] | 7.08<br>[5.90–8.85] | 3.23<br>[0.93–5.53] | 0.12<br>[0.11–0.13] | 0.15<br>[0.13–0.17] | 1.11<br>[0.84–1.38] | 15.62<br>[15.28–15.95] |
| C<sub>R2</sub> | 0.54<br>[0.43–0.65] | 6.52<br>[5.43–8.15] | 1.78<br>[0.60–2.96] | 0.10<br>[0.09–0.11] | 0.06<br>[0.05–0.07] | 1.90<br>[1.12–2.68] | 15.31<br>[14.02–16.60] |
| C<sub>R3</sub> | 0.39<br>[0.31–0.47] | 4.69<br>[3.90–5.86] | 1.71<br>[0.60–2.82] | 0.11<br>[0.10–0.12] | 0.07<br>[0.06–0.08] | 1.77<br>[1.50–2.04] | 15.33<br>[14.09–16.57] |
| C<sub>R4</sub> | 1.68<br>[1.57–1.78] | 0.60<br>[0.52–0.67] | 1.08<br>[0.73–1.42] | 0.33<br>[0.32–0.34] | 0.43<br>[0.40–0.47] | 0.63<br>[0.44–0.83] | 12.90<br>[12.58–13.23] |
| C<sub>R6</sub> | 1.49<br>[1.21–1.75] | 2.27<br>[2.00–2.55] | 1.21<br>[0.67–1.74] | 0.34<br>[0.33–0.35] | 0.50<br>[0.46–0.55] | 0.79<br>[0.64–0.93] | 14.97<br>[14.53–15.41] |
| C<sub>R7</sub> | 1.53<br>[1.24–1.80] | 2.65<br>[2.35–2.95] | 1.36<br>[0.91–1.81] | 0.32<br>[0.31–0.33] | 0.42<br>[0.40–0.46] | 0.74<br>[0.57–0.92] | 15.73<br>[15.12–16.33] |

### Prey morphology

To investigate whether the observed differences in defensive and competitive traits of *C. reinhardtii* strains were related to morphological differences among strains, we analyzed morphological traits in more detail. Strains exhibiting strong clumping morphologies (C_R1_, C_R2_ and C_R3_) displayed populations covering a larger spectrum of the PCA dimensional space (Fig. S4a) compared to strains exhibiting mostly single cell morphologies (C_R4_, C_R6_ and C_R7_). Clumping strains were positively associated to traits indicating large asymmetric cell aggregates (e.g., area and lobe count, Fig. S4b) and negatively associated to traits indicating small symmetric single cells (e.g., aspect ratio and roundness, Fig. S4b). This was confirmed by the negative correlations found between area and roundness for clumping strains (e.g., C_R2_: Pearson correlation, *r* = –0.43, *P* < 0.001) but not for non-clumping strains (e.g., C_R4_: Pearson correlation, *r* = 0.07, *P* < 0.001). Accordingly, significant differences among strains were found for area (Kruskal-Wallis, *χ^2^* = 14.37, *P* = 0.013) and roundness (Kruskal-Wallis, *χ^2^* = 8177.60, *P* < 0.001) indicating that clumping strains exhibited on average larger size and less symmetric shapes (Fig. S6). Overall, the presence or absence of clumping morphotypes coincided with the defensive and competitive traits of strains, meaning that strains having large cell clumps were classified as very defended and poorly competitive and vice versa (Fig. 3).

### Defense-competitiveness trade-off

We found strong negative relationships between anti-grazing defense and competitiveness traits independent of the trait used (Fig. 2). Significant relationships were found between attack rate, handling time or predator growth rate and prey maximum growth rate or nitrate affinity constant (Tab. 2). Except for C:N ratio, non-significant relationships between traits showed persistent relative positions in trait spaces with non-overlapping standard deviations. The more defended strains were consistently less competitive and vice versa. More defended strains (C_R1_, C_R2_ and C_R3_) were characterized by large-sized cell clumps (e.g., C_R1_: 323 ± 230 µm), resulting in low attack rates (e.g., C_R1_: 0.19 ± 0.04 10^-6^ mL sec^-1^) and high handling times (e.g., C_R1_: 4.64 ± 0.80 sec). Conversely, the more competitive strains (C_R4_, C_R6_ and C_R7_) were characterized by small-sized single cells (e.g., C_R4_: 108 ± 35 µm), resulting in high attack rates (e.g., C_R4_: 1.55 ± 0.11 10^-6^ mL sec^-1^) and low handling times (e.g., C_R4_: 0.56 ± 0.09 sec). The positions of strains in the trait space were similar for trait spaces constructed with prey maximum growth rates (Fig. 2a) or growth parameters related to nitrate, such as nitrate affinity constants (Fig. 2b) and half-saturation constants (Fig. 2c), as traits for competitiveness.

**Figure 2:**
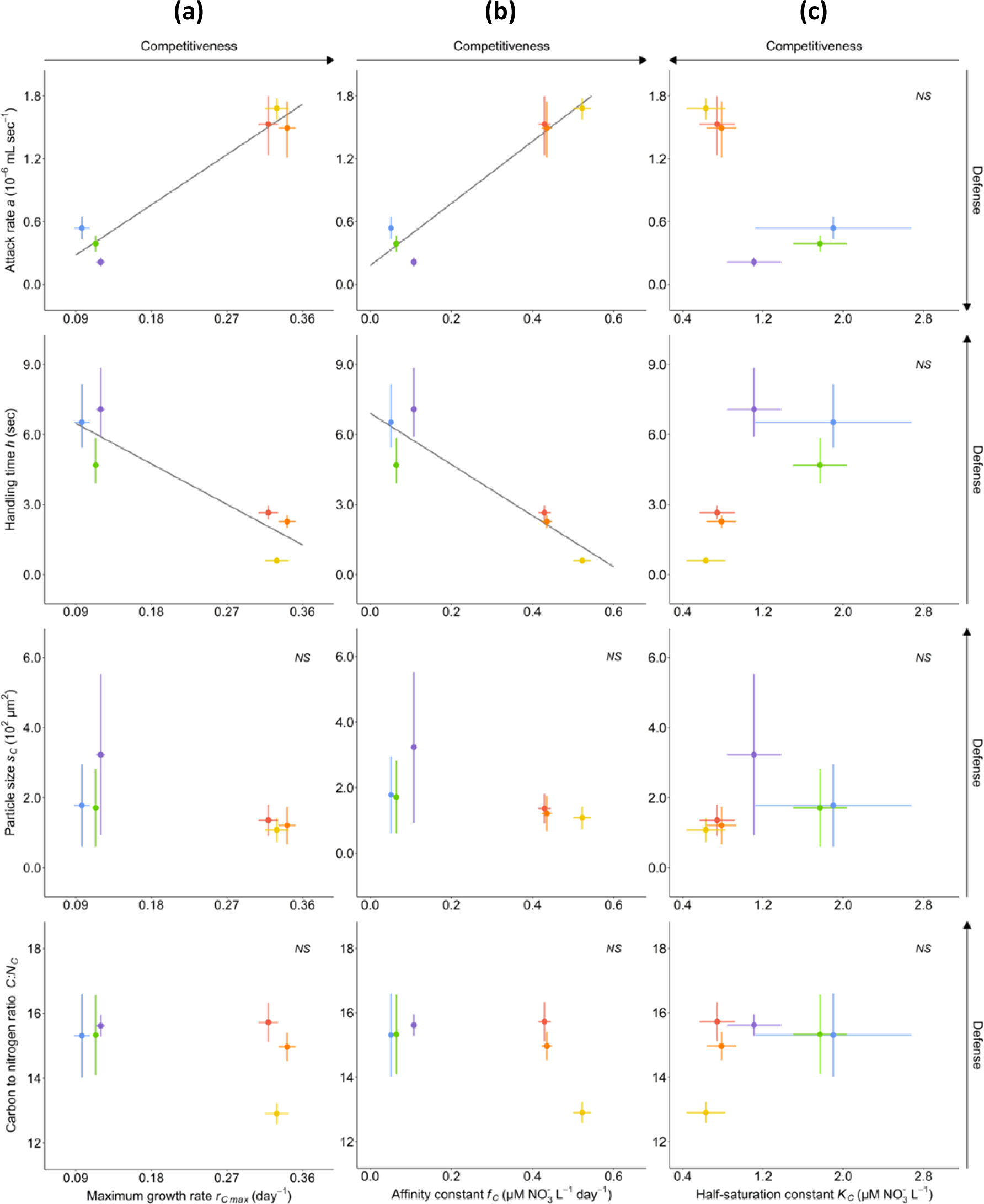
Defense-competitiveness trait spaces of *C. reinhardtii* strains. Shown are the positions for four anti-grazing defense traits (attack rate *a*, handling time *h*, prey particle size *s_C_* and carbon to nitrogen ratio *C:N_C_*) as a function of three competitiveness traits (prey maximum growth rate *r_Cmax_*, nitrate affinity constant *f_C_* and nitrate half-saturation constant *K_C_*). Trait estimates (dots) and standard deviations (horizontal and vertical bars) were represented pairwise for competitiveness traits on each plot: (a) prey maximum growth rate, (b) prey nitrate affinity constant and (c) prey nitrate half-saturation constant. Linear regression lines showing the significant correlations between anti-grazing defense and competitiveness traits were represented (grey lines) and non-significant correlation were indicated by a label (*NS*).

**Table 2:** Parameter estimates for parametric correlation tests performed between defense and competitiveness traits. Correlations were represented by regression lines on the different defense-competitiveness trait spaces (Fig. 2). Anti-grazing defense was represented by *B. calyciflorus* attack rate (*a* in 10^-6^ mL sec^-1^), handling time (*h* in sec), *C. reinhardtii* particle size (*s_C_* in 10^3^ µm^2^) and carbon to nitrogen ratio (*C:N_C_*). Competitiveness was represented by *C. reinhardtii* maximum growth rate (*r_Cmax_* in day^-1^), affinity constant (*f_C_* in µmol NO_3_^-^ L^-1^ day^-1^) and half-saturation constant (*K_C_* in µmol NO_3_^-^ L^-1^) for nitrate. Significance was estimated with a Pearson correlation test (*t*, *df* and *P* values).

| Defense trait | Competitiveness trait | $r$ | $t$ | $df$ | $P$ |
| --- | --- | --- | --- | --- | --- |
| $a$ | $r_{Cmax}$ | 0.97 | 7.90 | 4 | 0.001 |
| $h$ | | -0.91 | -4.41 | 4 | 0.012 |
| $s_C$ | | -0.69 | -1.89 | 4 | 0.132 |
| $C:N_C$ | | -0.47 | -1.06 | 4 | 0.350 |
| $a$ | $f_C$ | 0.97 | 7.85 | 4 | 0.001 |
| $h$ | | -0.93 | -4.86 | 4 | 0.008 |
| $s_C$ | | -0.66 | -1.76 | 4 | 0.154 |
| $C:N_C$ | | -0.58 | -1.42 | 4 | 0.229 |
| $a$ | $K_C$ | -0.79 | -2.59 | 4 | 0.061 |
| $h$ | | 0.73 | 2.14 | 4 | 0.099 |
| $s_C$ | | 0.30 | 0.63 | 4 | 0.566 |
| $C:N_C$ | | 0.42 | 0.93 | 4 | 0.404 |

### Prey and predator fitness

The defensive and competitive traits of *C. reinhardtii* strains had an impact on the population growth of the prey and the predator (Fig. 3). The comparison of prey growth models revealed that *C. reinhardtii* strains had a logistic growth (Tab. S4 and Fig. S1b). More defended strains had lower growth rates and asymptotic densities under non-limiting nitrate conditions compared to less defended strains (Tab. S6). The less defended strain C_R4_ showed a rapid increase followed by a decrease in cell density probably attributed to depletion of nitrate and/or shading at high densities (Fig; 3a). The comparison of predator growth models revealed that *B. calyciflorus* had a logistic growth in the presence of the less defended strains C_R6_ and C_R7_ but a logistic growth followed by a decline in the presence of the more defended strains C_R1_, C_R2_ and C_R3_ and the less defended strain C_R4_ (Fig. 3b; Tab. S5 and Fig. S1c). More defended strains were associated with lower predator growth rates (e.g., C_R1_: −0.05 ± 0.18 day^-1^) and asymptotic densities while less defended strains were associated with higher predator growth rates (e.g., C_R4_: 2.33 ± 0.27 day^-1^) and asymptotic densities (Tab. S6). The less defended strain C_R4_ showed a rapid increase in rotifer density followed by a decrease probably attributed to a strong prey depletion.

**Figure 3:**
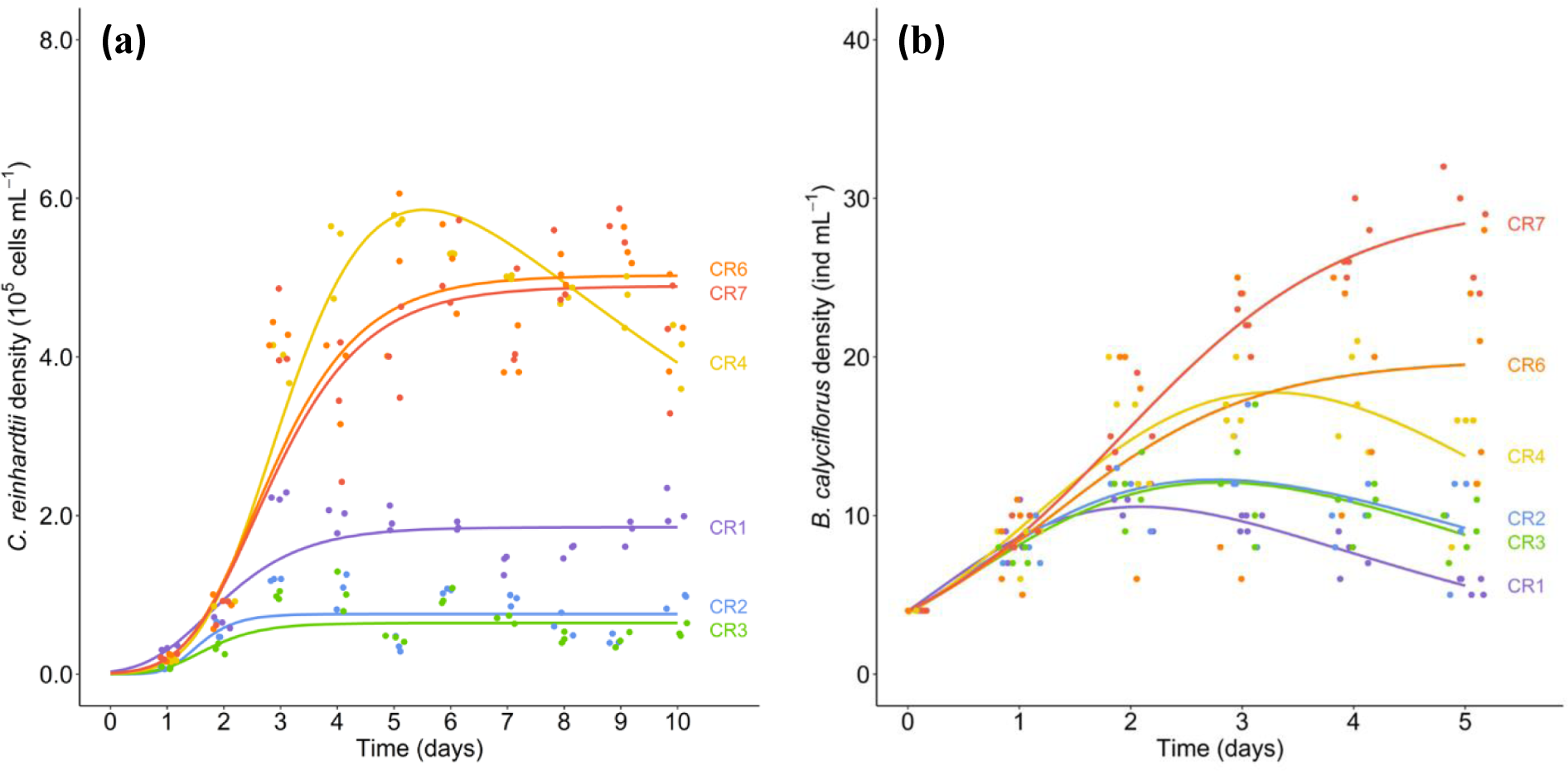
Fitness *C. reinhardtii* strains and *B. calyciflorus*. (a) Growth curves of *C. reinhardtii* strains shown as cell densities over time for a high nitrate concentration (100 µmol NO_3_^-^ L^-1^). Predicted growth curves (lines) were estimated from the 3 replicates for cell densities (dots). (b) Growth curves of *B. calyciflorus* feeding on *C. reinhardtii* strains shown as rotifer densities over time. Predicted growth curves (lines) were estimated from the 5 replicates of rotifer densities (dots). Predicted curves with lower and upper boundaries using standard errors are shown in the *Supplementary Materials* (Fig. S2a and S2b).

## Discussion

Defense-competitiveness trade-off relationships are often observed in plankton organisms and can be expressed with multiple traits (Pančić and Kiørboe 2018). Determining the relevant traits describing these trade-offs is essential to predict the outcome of trophic interactions (Cadier et al. 2019). Comparing multiple anti-grazing defense and competitiveness traits, we found substantial variation among strains of *C. reinhardtii* and overall negative relationships between these traits. The negative relationships between defense and competitiveness traits suggests a colimitation in energy investment and/or an evolutionary constrain between survival and reproduction (Stearns 1989). This observation also shows that multiple morphological and trophic traits of prey and predator can be suitable proxies to estimate trade-off relationships and a mechanistic link between prey structural traits and predator ingestion and reproduction traits (Litchman et al. 2013). The differences in defense and competitiveness traits and their relationships translated into different fitness consequences for prey and predator and thus their interaction strength. We found lower population sizes of *B. calyciflorus* in the presence of more defended *C. reinhardtii* strains, confirming that prey defensive and competitive traits, predator-prey trophic interactions and predator reproduction are closely connected (Litchman et al. 2013). Our study confirms previous observations comparing trait variation between species or within species of higher taxonomic groups (Litchman et al. 2013; Ehrlich et al. 2020) and improves the understanding of planktonic communities (Litchman et al. 2007; 2010) by demonstrating the mechanistic links between traits, trade-offs and fitness at the intraspecific level.

Our data suggest a mechanistic relationship at the intraspecific level linking prey morphological traits to predator trophic traits. *C. reinhardtii* strains classified as defended against ingestion by the predator were growing as cell groups bound by a viscous extracellular matrix. These strains were associated with lower predator functional responses which were mainly the consequence of the gape-size limitation of the predator for ingesting cell groups within an extracellular matrix (Reynolds 2006). Lower ingestion rates of cell colonies could also indicate longer manipulation and digestion times for the predator (Verschoor et al. 2007), and/or a resistance to digestion during gut passage (DeMott et al. 2010). Predator traits such as attack rate and handling time are reliable indicators of prey defense (Jeschke and Tollrian 2000) and other traits related to ingestion such as digestion time and conversion efficiency (Jeschke et al. 2002) should be further explored. Prey morphological traits can also provide relevant estimates of prey defense (e.g., Becks et al. 2010; Ellner and Becks 2011). Prey size and shape is a particularly pertinent trait as it is directly associated with gape-size limitation for the predator. Some caution is still required with prey size and shape as other morphological traits (e.g., spines) or other physiological (e.g., toxins) and behavioral (e.g., motility) traits can contribute to prey defense (Pančić and Kiørboe 2018; Lürling 2021) and the relative contribution of these traits to prey defense is often unclear (Van Donk et al. 2011).

Our data also suggest a mechanistic relationship at the intraspecific level linking prey morphological traits to prey growth traits. *C. reinhardtii* strains classified as competitive were associated with higher growth rates and nitrate affinities but lower nitrate half-saturation constants, which indicated enhanced abilities for nutrient acquisition or conversion to reproduction (Litchman et al. 2007; 2010; Halsey and Jones 2015; Ward et al. 2017). Competitive abilities of phytoplankton species mainly depend on surface-area constrains in terms of resource acquisition and requirements for basal maintenance and functions contributing to fitness. Single cells tend to have higher nutrient acquisition rates due to a more efficient diffusion of nutrients (Yoshiyama and Klausmeier 2008) while palmelloid colonies are more limited by nutrient and light competition among colonial cells (Raven and Kubler 2002), despite larger nutrient storage capacities (Litchman et al. 2009). Prey growth rate and nutrient acquisition rate are reliable indicators of prey competitiveness and show a relation between the processes of nutrient absorption, assimilation, and conversion to reproduction (Litchman et al. 2007). However, complex interplays between processes may complicate predictions as prey competitiveness can also be driven by other regulating factors such as the acquisition of colimiting resources (Klausmeier et al. 2004) or trade-offs between nutrient and light acquisition (Strzepek and Harrison 2004).

The intraspecific covariation in defense and competitiveness traits of phytoplankton have consequences on the growth of zooplankton. Prey defense, including morphological and physiological traits, and prey stoichiometry have been shown to affect food quantity and quality (Bi and Sommer 2020), which prevents the predator from meeting its energetic requirements and thus reduces its survival and reproduction (Sommer 2008). Prey stoichiometry is often used as a good predictor of prey defense and competitiveness traits and thus predator fitness (Sterner and Elser 2002) as C:N (and C:P) ratio increases due to the high carbon requirements when cells form colonies. Conversely, C:N (and C:P) ratio decreases due to the high nitrogen and phosphorus requirements of cell replication (Branco et al. 2020). Such differences in prey stoichiometry can impact predator growth rate (Branco et al. 2010; 2018; Mandal et al. 2018) but we did not observe differences in C:N ratios among *C. reinhardtii* strains. Whether this is due to the precise growth conditions (e.g., nutrient concentration or light intensity) or indicates a difference between intraspecific and interspecific variation should be further studied.

We used 6 strains of *C. reinhardtii* with a known recent evolutionary history (Bernardes et al. 2021) to examine defense-competitiveness trade-off relationships. The isolates came from an evolution experiment in which either defense or competitiveness was selected for and a general trade-off between survival against predation and replication was observed across a large range of isolates (Bernardes et al. 2021). Previous approaches consisted in split-family experiments in which clones were evolved in different environments and thus this could affect the fitness of clones in an opposite way leading to no or positive correlations between traits (Fry 1993; 1996). The experimental evolution approach has been proven to be more suitable for detecting trade-offs than the split-family approach (Kassen 2002; Fry 2003; Fuller et al. 2005). Selection experiments can also be used to determine the role of phenotypic plasticity (Scheiner 2002) and environmental heterogeneity (Kassen 2002) versus genetic adaptation for the maintenance of trait variation (Roff and Fairbairn 2007). Overall, this approach allows testing whether 2 traits can be optimized concurrently or whether optimization of one trait leads to a reduction in the other trait.

Investigating intraspecific relationships between defensive and competitiveness traits is fundamental to understand fitness strategies of interacting organisms (Ehrlich et al. 2017; Bartlett et al. 2018) and coexistence between resources and consumers (Lichstein et al. 2007; Petry et al. 2018; Ehrlich et al. 2020). Our study demonstrates intraspecific defense-competitiveness trade-offs via negative relationships estimated over a range of traits measured on both prey and predator. Assessing the magnitude and the shape of intraspecific trade-offs can be decisive as empirical and theoretical studies showed the significant role of trade-offs for consumer-resource dynamics (e.g., Yoshida et al. 2003; Becks et al. 2010; Cortez and Ellner 2010; Kasada et al. 2014; Ehrlich et al. 2017). Knowledge on the underlying traits involved in trade-offs and how these trade-offs translate into fitness of interacting organisms at different trophic levels can provide key insights into the transfers of energy and matter in food webs (Våge et al. 2014; 2018; Cadier et al. 2019) and the response of populations to environmental perturbations (Jessup and Bohannan 2008; Frickel et al. 2017; Theodosiou et al. 2019). Intraspecific trade-offs are for instance a prerequisite for eco-evolutionary feedback dynamics, which can contribute to the maintenance of trait variation (e.g., Becks et al. 2010; Kasada et al. 2014) and in turn allow the persistence of populations (Yamamichi and Miner 2015; Hermann and Becks 2022). Investigating intraspecific trade-offs thus offers a promising avenue for assessing trait variation within population and predict the strength of trophic interactions, population dynamics and ecosystem processes.

## Acknowledgements

We are thankful to U. Gaedke and Z. Zhang for comments on an earlier version the manuscript, to E. Ehrlich for the advice on the data analysis, and to R. Hermann and R. Lambrecht for the theoretical and practical contributions to the experiments.

## Author contributions

TR and LB compiled the first theoretical ideas. TR and LB designed the experiments. TR conducted the laboratory experiments and statistical analyses with the assistance of LB. TR wrote the initial drafts of the manuscript. TR and LB wrote the final version of the manuscript.

