## Supplementary Materials for "Multiple trade-offs between defense and competitiveness traits in a planktonic predator-prey system"

### Appendix 1: images acquisition and treatment using a high contrast imaging microscope

Image acquisitions were performed using an integrated cellular imaging and analysis microscope designed for screening of fluorescent biological samples on microplates (ImageXpress® Micro 4 High Content Imaging System). Acquisitions were made using a 96 well plate design with 21 sites per well (grid: 5×5 sites) covering 26.25 $\mu m^2$ (surface: 77.52 %) under a 10X magnification. A filter (Cyt5) selecting for red fluorescence (644−712 nm) in bright light emitted by a laser was used to visualize excited chlorophyll in cells. Cell images were obtained with an exposure time of 200 msec and a focus offset of −2.78 µm. To capture cells that did not sink to the bottom of the well but were floating on top of the water column, we performed a second acquisition using a vertical recording (*Z series* function) of 11 images in heights ranging from 3200 to 4200 µm (step: 100 µm) with an automatic shading correction and selecting the sum projection of images. Image analyses to count cells were performed using a related software (MetaXpress® High Content Image Acquisition and Analysis Software). A custom module was created (ImageXpress® Module Cell Counting) and applied on images to facilitate cell counting at both bottom and top of wells. For the well bottom, a *top hat* function (i.e., finds small bright spots in objects based on a filter size and shape) was used to accentuate intensity of circular cells of maximum 20 µm. A *find round object* function (i.e., identifies small circular objects using size and fluorescence intensity) was then used to count cells ranging from 4 to 8 µm having an intensity of 500 msec above the dark background. Cell densities were estimated from cell counts as $D = N_C \times N_S \times D_F \times C_F$, where $D$ is the cell density (cells $mL^{-1}$), $N_S$ is the number of images recorded per well, $D_F$ is the dilution factor to convert the cell density per mL and $C_F$ is the coverage factor to rescale the cell density covered by the images to the total surface of the well.

### Appendix 2: images acquisition and treatment using an imaging flow cytometer

Image acquisitions were performed using an integrated multichannel cellular imaging and analysis flow cytometer designed for imaging of fluorescent biological samples (Amnis® ImageStreamX Mk II). Acquisitions were made by loading 1.5 mL tubes containing 200 µL in the sample reader at low flow speed (200 cells sec$^{-1}$) under 40X magnification and an automatically adjusted focus offset. Illumination settings for the 4 lasers of the machine were calibrated as follows: blue fluorescence (405 nm) at 1 mW, green fluorescence (488 nm) at 1 mW, red fluorescence (642 nm) at 2 mW and side scatter fluorescence (785 nm) at 0.1 mW. Image analyses were performed using a related analysis software (Amnis® IDEAS). Images visualization was made for 12 image channels characterized by differences in fluorescence and channel 5 (Cyt5) corresponding to a red fluorescence (642 nm) was selected for the gating of images. A gating of images was made by scatter plotting the features Area as a function of Intensity both calculated on image channel 5 and by drawing a rectangular region (area = 10−500 µm; intensity = $10^1$−$10^6$ pixels) to create a gated population. Gating images was excluding small non-biological plastic speed beads (3 µm) from the image collection. Image acquisitions were made for the image channel 5 (5000 images per channel) and 9 features per image were calculated.

### Appendix 3: correction of predator functional responses for prey density depletion

The functional response models describe how predator ingestion rate changes with prey density in the environment. However, it is difficult to maintain prey density constant over the time of feeding experiments and this affects the estimation of the funtionnal parameters (*a*, *h* and *c*). Because *C. reinhardtii* densities ($C_R$) were not maintained constant over the time of our feeding experiment, we corrected *B. calyciflorus* ingestion rate ($i_R$) for each strain to account for prey depletion over time following Rosembaum and Rall (2018). We used ordinary differential equations modeling the dynamics of prey density (Eq. S1) and predator ingestion rate (Eq. S2) over time using Holling type II or Holling type III functional response:

$$\frac{dC_{Ri}}{dt} = C_{Ri} - \frac{a_i {C_{Ri}}^c}{1 + a_i h_i {C_{Ri}}^c} B_C \qquad (\text{Eq. S1})$$

$$\frac{di_{Ri}}{dt} = \frac{a_i {C_{Ri}}^c}{1 + a_i h_i {C_{Ri}}^c} B_C \qquad (\text{Eq. S2})$$

where $a_i$ is the attack rate of the predator on strain *i* ($10^{-6}$ mL $sec^{-1}$), $h_i$ is the handling time of the predator on strain *i* (sec), $c_i$ is the prey density exponent of the predator on strain *i*, $C_{Ri}$ is the cell density of strain *i* (cells $mL^{-1}$) and $B_C$ is the predator density (rotifers). We computed a log-likelihood function to estimate the functional parameters assuming a normal distribution for each log-transformed prey density using the *dnorm* function from the 'stats' R package. We fitted the log-likelihood function with a maximum likelihood estimation providing the mean and the standard deviation estimates of functional parameters using the *mle2* function from the 'bbmle' R package. We then solved the ordinary different equations system parametrized with functional parameters estimates for the time (8 hours) and the predator density (4 rotifers) of our feeding experiment using the *lsoda* function from the 'deSolve' R package. More information is available in the Supplementary Materials of Rosenbaum and Rall (2018).

**Table S1:** List of the 6 *C. reinhardtii* strains used in the experiments. Strains stem from different experimental lineages that were selected from naïve monoclonal ancestors during thousands of generations (Bernandes et al. 2021). Strains name and code originate from the Chlamydomonas Resource Center (https://www.chlamycollection.org).

| *Strain* | | *Code* | *Description* |
|---|---|---|---|
| $C_{R1}$ | 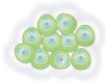 | cc1883 | Large cell clumps |
| $C_{R2}$ | 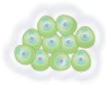 | cc3754 | Medium cell clumps |
| $C_{R3}$ | 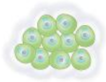 | cc3056 | Medium cell clumps |
| $C_{R4}$ | 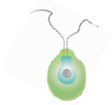 | cc2854 | Single and motile cells |
| $C_{R6}$ | 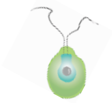 | cc1144 | Single and non-motile cells |
| $C_{R7}$ | 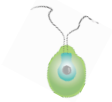 | cc2698 | Single and non-motile cells |

**Table S2:** List of the 9 morphological features used for the principal components analysis. Features are calculated on the image collections acquired with a flow cytometer (see *Appendix 2*) and estimated on a grid of 2000 pixels (25 $\mu m^2$ per pixel) covering images of individual cells.

| *Feature* | *Category* | *Description* |
|---|---|---|
| Area | Size | Surface covered by the cell ($\mu m^2$) |
| Aspect ratio | Shape | Ratio of the minor axis on the major axis |
| Roundness | Shape | Degree of deviation from the circle of the cell area |
| Diameter | Size | Diameter of the circle of the cell area (µm) |
| Elongation | Shape | Ratio of the height on the width of the cell |
| Length | Size | Length of the longest part of the cell (µm) |
| Lobe count | Shape | Number of lobes within the cell based on symmetry |
| Perimeter | Size | Length of the boundary of the cell area (µm) |
| Width | Size | Length of the shorter side of a bounding rectangle (µm) |

**Table S3:** Ingestion models used to calculate functional response curves of *B. calyciflorus* on *C. reinhardtii* strains (Fig. 1a). The table shows model formulations expressing ingestion rate ($i_R$ in cells $sec^{-1}$ $ind^{-1}$) as a function of the initial cell density intercept ($C_{R0}$ in $10^6$ cells $mL^{-1}$), the attack rate ($a$ in $10^{-6}$ mL $sec^{-1}$), the handling time ($h$ in sec), the prey density exponents ($c$ and $d$) and the prey density ($C_R$ in $10^6$ cells $mL^{-1}$). Functional response models were selected per strain based on the lowest AIC values. Non-fitting ingestion models were indicated by a horizontal bar (−) and the choice of models was indicated (×).

| *Model* | *Formulation* | *Strain* | *AIC* | *Fitted* |
|---|---|---|---|---|
| Holling I | $i_R = C_{R0} + aC_R$ | $C_{R1}$ | −72.28 | · |
| | | $C_{R2}$ | −61.97 | · |
| | | $C_{R3}$ | −72.00 | · |
| | | $C_{R4}$ | −18.40 | · |
| | | $C_{R6}$ | −30.67 | · |
| | | $C_{R7}$ | −36.54 | · |
| Holling II | $i_R = \frac{aC_R}{1 + ahC_R}$ | $C_{R1}$ | −78.42 | · |
| | | $C_{R2}$ | −67.97 | × |
| | | $C_{R3}$ | −78.63 | × |
| | | $C_{R4}$ | −32.78 | · |
| | | $C_{R6}$ | −42.50 | × |
| | | $C_{R7}$ | −39.78 | × |
| Holling III | $i_R = \frac{a{C_R}^c}{1 + ah{C_R}^c}$ | $C_{R1}$ | −82.65 | × |
| | | $C_{R2}$ | −65.97 | · |
| | | $C_{R3}$ | −76.64 | · |
| | | $C_{R4}$ | −41.30 | × |
| | | $C_{R6}$ | −40.73 | · |
| | | $C_{R7}$ | −39.35 | · |
| Ivlev II | $i_R = \frac{(1 - e^{-dC_R})}{h}$ | $C_{R1}$ | −79.07 | · |
| | | $C_{R2}$ | −67.00 | · |
| | | $C_{R3}$ | −78.50 | · |
| | | $C_{R4}$ | −35.33 | · |
| | | $C_{R6}$ | −41.95 | · |
| | | $C_{R7}$ | −37.92 | · |

**Table S4:** Growth models used to calculate growth curves of *C. reinhardtii* strains (Fig. 1b). The table shows model formulation expressing cell density ($C_R$ in $10^6$ cells $mL^{-1}$) as a function of the initial cell density ($C_{R0}$ in $10^6$ cells $mL^{-1}$), the intrinsic growth rate ($r_C$ in $day^{-1}$), the asymptotic cell density ($c_C$ in $10^6$ cells $mL^{-1}$), the decreasing slope following maximum cell density ($m_C$ in $day^{-1}$), the time ($t$ in day) and the time of maximum intrinsic growth rate ($t_m$ in day). Growth rate models were selected per strain based on the lowest AIC values. Non-fitting growth models were indicated by a horizontal bar (−) and the choice of models was indicated (×).

| *Model* | *Formulation* | *Strain* | *AIC* | *Fitted* |
|---|---|---|---|---|
| Linear | $C_R = C_{R0} + r_C t$ | CR1 | 18.84 | · |
| | | CR2 | 27.88 | · |
| | | CR3 | 25.76 | · |
| | | CR4 | 30.25 | · |
| | | CR6 | 27.43 | · |
| | | CR7 | 26.63 | · |
| Exponential | $C_R = C_{R0} + e^{r_C t}$ | CR1 | 18.98 | · |
| | | CR2 | 27.94 | · |
| | | CR3 | 25.81 | · |
| | | CR4 | 30.62 | · |
| | | CR6 | 27.84 | · |
| | | CR7 | 27.08 | · |
| Logistic | $C_R = \frac{C_{R0} c_C}{C_{R0} + (c_C - C_{R0}) e^{-r_C t}}$ | CR1 | 5.94 | × |
| | | CR2 | 16.71 | × |
| | | CR3 | 15.01 | × |
| | | CR4 | 1.78 | · |
| | | CR6 | 4.64 | × |
| | | CR7 | 4.84 | × |
| Gompertz | $C_R = c_C \left(\frac{C_{R0}}{c_C}\right)^{e^{-r_C t}}$ | CR1 | 6.13 | · |
| | | CR2 | − | · |
| | | CR3 | − | · |
| | | CR4 | 2.83 | · |
| | | CR6 | 5.24 | · |
| | | CR7 | 5.25 | · |
| Mortality | $C_R = c_C e^{m_C (t - t_m) - \left(\frac{m_C}{r_C}\right)\left(1 - e^{-r_C (t - t_m)}\right)}$ | CR1 | − | · |
| | | CR2 | − | · |
| | | CR3 | − | · |
| | | CR4 | −4.49 | × |
| | | CR6 | − | · |
| | | CR7 | − | · |

**Table S5:** Growth models used to calculate growth curves of *B. calyciflorus* (Fig. 1d). The table shows model formulation expressing rotifer density ($B_C$ in rotifers mL$^{-1}$) as a function of the initial rotifer density ($B_{C0}$ in rotifers mL$^{-1}$), the intrinsic growth rate ($r_B$ in day$^{-1}$), the asymptotic rotifer density ($c_B$ in rotifers mL$^{-1}$), the decreasing slope following maximum rotifer density ($m_B$ in day$^{-1}$), the time (*t* in day) and the time of maximum intrinsic growth rate ($t_m$ in day). Growth rate models were selected per strain based on the lowest AIC values. Non-fitting growth models were indicated by a horizontal bar (−) and the choice of models was indicated (×).

| *Model* | *Formulation* | *Strain* | *AIC* | *Fitted* |
|---|---|---|---|---|
| Linear | $B_C = B_{C0} + r_B t$ | $C_{R1}$ | 32.82 | · |
| | | $C_{R2}$ | 32.68 | · |
| | | $C_{R3}$ | 32.54 | · |
| | | $C_{R4}$ | 36.74 | · |
| | | $C_{R6}$ | 27.86 | · |
| | | $C_{R7}$ | 28.26 | · |
| Exponential | $B_C = B_{C0} + e^{r_B t}$ | $C_{R1}$ | 32.83 | · |
| | | $C_{R2}$ | 33.24 | · |
| | | $C_{R3}$ | 33.07 | · |
| | | $C_{R4}$ | 38.06 | · |
| | | $C_{R6}$ | 33.14 | · |
| | | $C_{R7}$ | 36.56 | · |
| Logistic | $B_C = \frac{B_{C0} c_B}{B_{C0} + (c_B - B_{C0}) e^{-r_B t}}$ | $C_{R1}$ | – | · |
| | | $C_{R2}$ | 26.19 | · |
| | | $C_{R3}$ | 26.39 | · |
| | | $C_{R4}$ | 29.15 | · |
| | | $C_{R6}$ | 8.66 | × |
| | | $C_{R7}$ | 12.79 | × |
| Mortality | $B_C = c_B e^{m_B(t-t_m) - \left(\frac{m_B}{r_B}\right)\left(1-e^{-r_B(t-t_m)}\right)}$ | $C_{R1}$ | 14.01 | × |
| | | $C_{R2}$ | 17.99 | × |
| | | $C_{R3}$ | 12.01 | × |
| | | $C_{R4}$ | 20.55 | × |
| | | $C_{R6}$ | – | · |
| | | $C_{R7}$ | – | · |

**Table S6:** Estimates of fitness proxies [± standard deviation] for the 6 different *C. reinhardtii* strains and *B. calyciflorus*: *C. reinhardtii* growth rate ($r_C$ in day$^{-1}$), asymptotic cell density ($c_C$ in $10^5$ cells mL$^{-1}$), *B. calyciflorus* growth rate ($r_B$ in day$^{-1}$) and asymptotic rotifer density ($c_C$ in rotifers mL$^{-1}$). Strains exhibited different morphologies in terms of cell clumping: clumping strains were $C_{R1}$ (large clumps [10−30 cells]), $C_{R2}$ (medium clumps [5−10 cells]) and $C_{R3}$ (medium clumps [5−10 cells]) while non-clumping strains were $C_{R4}$ (small single cells), $C_{R6}$ (small single cells) and $C_{R7}$ (small single cells).

| *Strain* | | $r_C$ | $c_C$ | $r_B$ | $c_B$ |
|---|---|---|---|---|---|
| $C_{R1}$ | 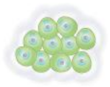 | 0.75 [0.71−0.80] | 1.85 [1.48−2.23] | −0.05 [−0.23−0.13] | 10.55 [10.54−10.57] |
| $C_{R2}$ | 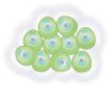 | 0.97 [0.86−1.09] | 0.76 [0.61−0.91] | 0.96 [0.78−1.14] | 12.26 [12.10−13.40] |
| $C_{R3}$ | 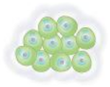 | 0.89 [0.81−0.97] | 0.64 [0.52−0.77] | 0.94 [0.76−1.12] | 12.09 [11.95−12.45] |
| $C_{R4}$ | 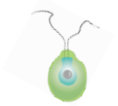 | 1.48 [1.42−1.54] | 5.86 [5.83−6.96] | 2.33 [2.08−2.58] | 17.76 [17.24−19.51] |
| $C_{R6}$ | 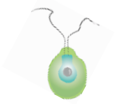 | 1.30 [1.24−1.36] | 5.02 [4.02−6.03] | 3.33 [3.20−3.46] | 19.51 [19.35−19.68] |
| $C_{R7}$ | 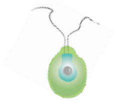 | 1.27 [1.22−1.33] | 4.89 [3.91−5.87] | 5.47 [5.35−5.59] | 28.41 [27.50−29.32] |

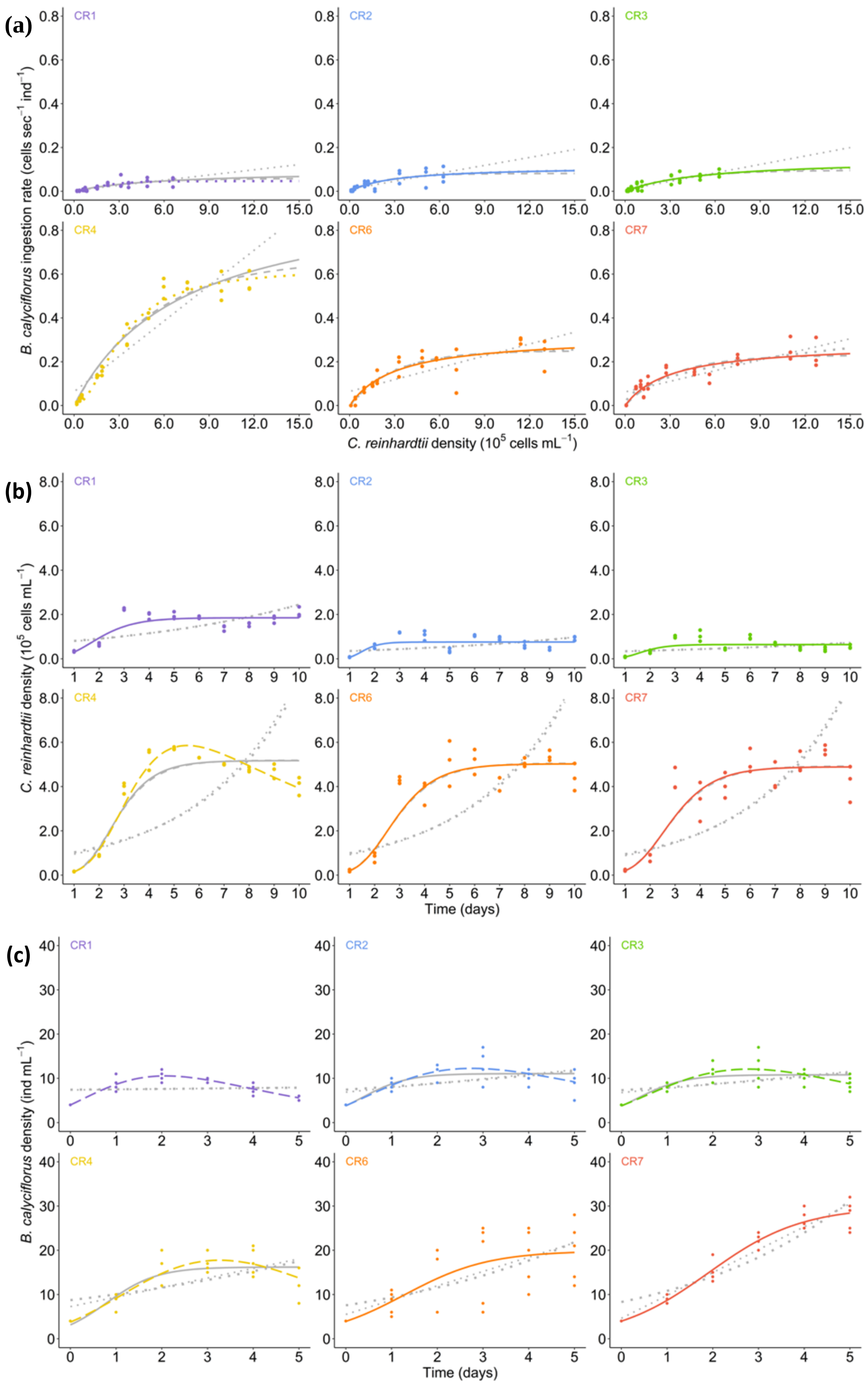
(a)
(b)
(c)
CR1
CR2
CR3
CR4
CR6
CR7
B. calyciflorus ingestion rate (cells sec−1 ind−1)
C. reinhardtii density (10^5 cells mL−1)
C. reinhardtii density (10^5 cells mL−1)
Time (days)
B. calyciflorus density (ind mL−1)
Time (days)

**Figure S1:** (a) Comparison of models for functional responses of *B. calyciflorus* on *C. reinhardtii* strains. Predicted functional response curves (lines) were estimated from the 3 replicates for ingestion rates (dots). Ingestion models: Holling type I (small dotted line), Holling type II (solid line), Holling type III (large dotted line) and Ivlev (short dashed line). (b) Comparison of models for growth curves of *C. reinhardtii* strains. Predicted growth curves (lines) were estimated from the 3 replicates for cell densities (dots). Growth models: Linear (small dotted line), Exponential (large dotted line), Logistic (solid line), Gompertz (short dashed line) and Mortality (long dashed line). (c) Comparison of models for growth curves of *B. calyciflorus* on *C. reinhardtii* strains. Predicted growth curves (lines) were estimated from the 3 replicates for rotifer densities (dots). Growth models: Linear (small dotted line), Exponential (large dotted line), Logistic (solid line) and Mortality (long dashed line). Model selection was indicated by the line color transparency: selected models (colored line) and non-selected models (grey line).

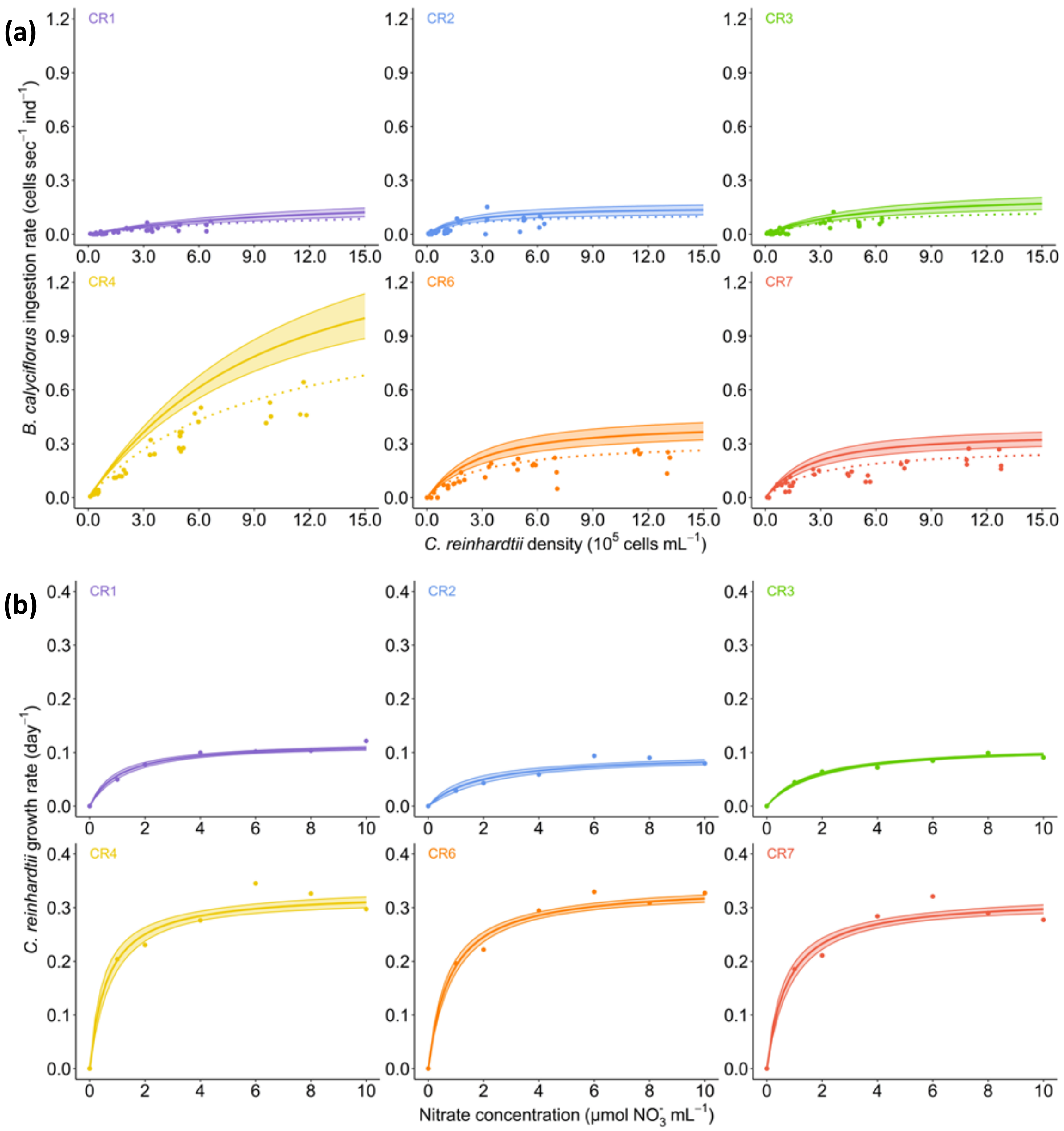


**Figure S2:** (a) Functional responses of *B. calyciflorus* showing ingestion rate as a function of cell density of *C. reinhardtii* strains. Predicted functional response curves (lines) were estimated with lower and upper boundaries using standard errors from the 3 replicates for ingestion rates (dots) for raw data (dotted lines) and prey density-corrected data (solid lines). (b) Growth rate of *C. reinhardtii* strains over low nitrate concentrations (1−10 µmol $NO_3^-$ $L^{-1}$). Predicted curves were estimated with lower and upper boundaries using standard errors from the growth rate at each nitrate concentration (dots).

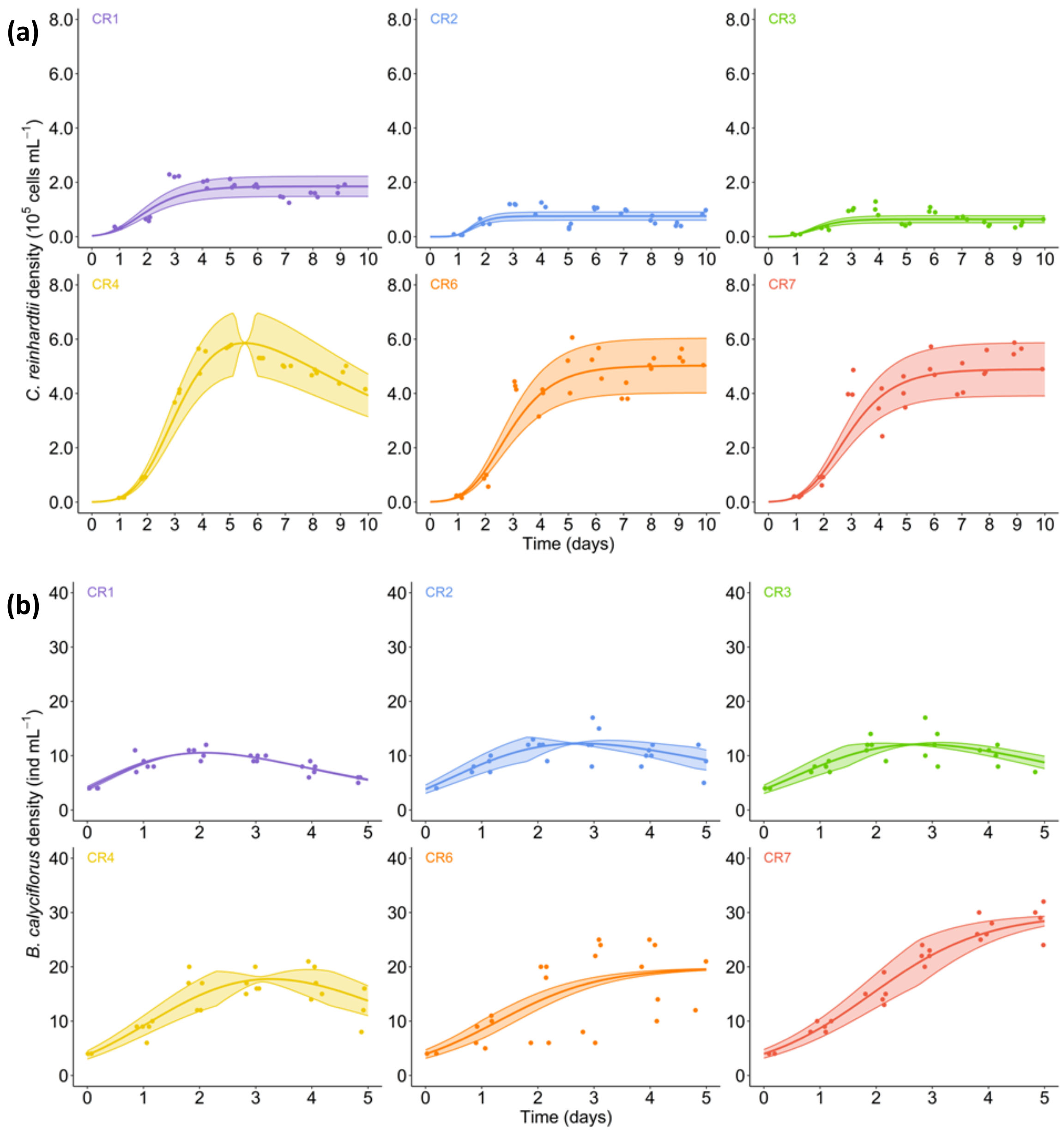


**Figure S3:** (a) Growth curves of *C. reinhardtii* strains showing cell density as a function of time for an intermediate nitrate concentration (100 µmol $NO_3^-$ $L^{-1}$). Predicted growth curves (lines) were represented with lower and upper boundaries using standard errors from the 3 replicates for cell densities (dots). (c) Growth curves of *B. calyciflorus* feeding on *C. reinhardtii* strains showing rotifer density as a function of time. Predicted growth curves (lines) were represented with lower and upper boundaries using standard errors from the 5 replicates for rotifer densities (dots).

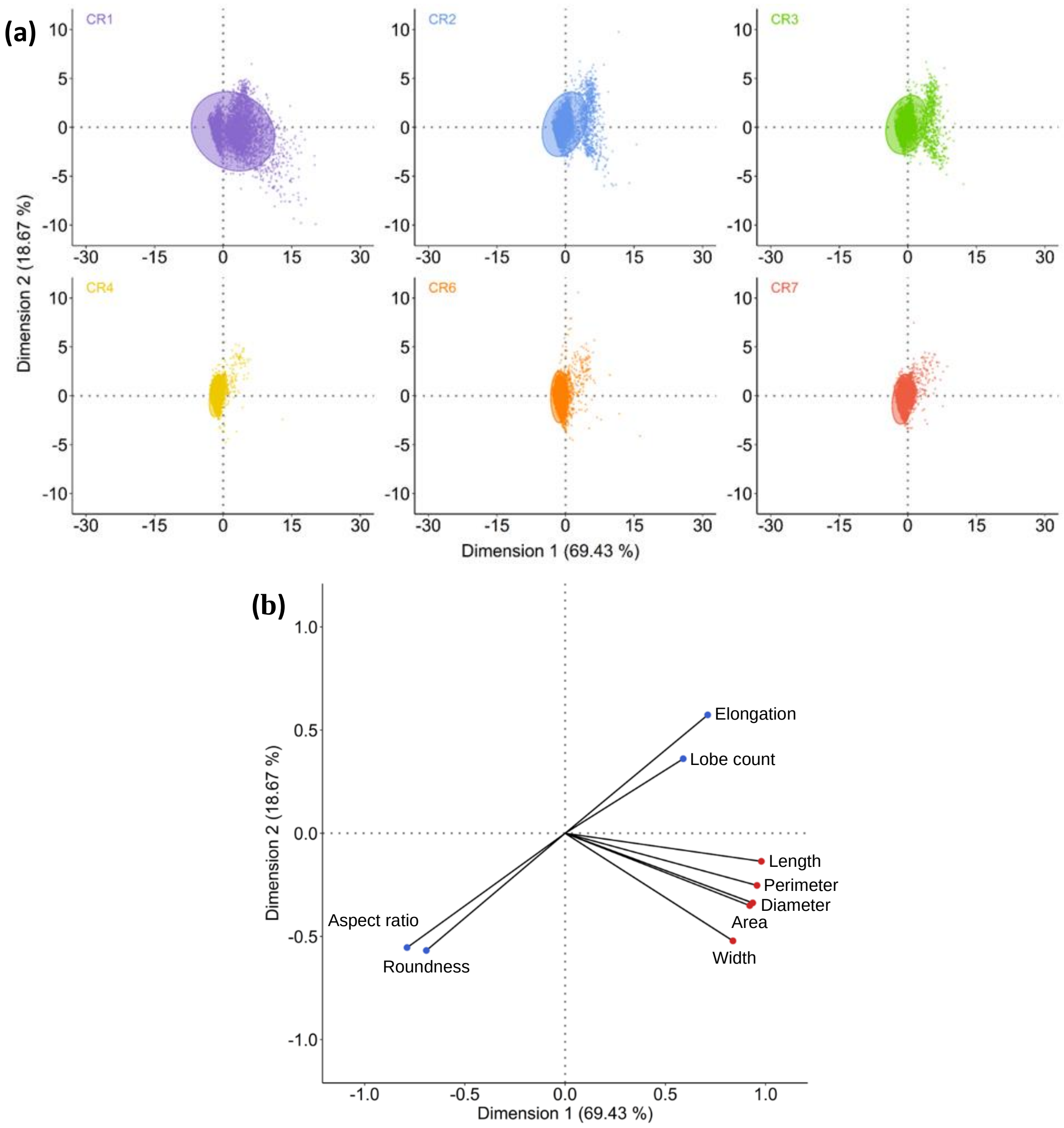


**Figure S4:** Representation of the principal components analysis (PCA) showing (a) the position of individual cell observations for populations of *C. reinhardtii* strains (n = 5000) and (b) the positions of morphological features (n = 9) along the 2 main dimensions explaining most of the variance ($D_1$ = 46.5% and $D_2$ = 19.9%). Ellipses group 90% of the cell observations for each strain population within strains. Features represented here were selected based on the highest contributions to the 2 main dimensions according to the PCA and were divided in 2 categories: cell size features (red dots) and cell shape features (blue dots).

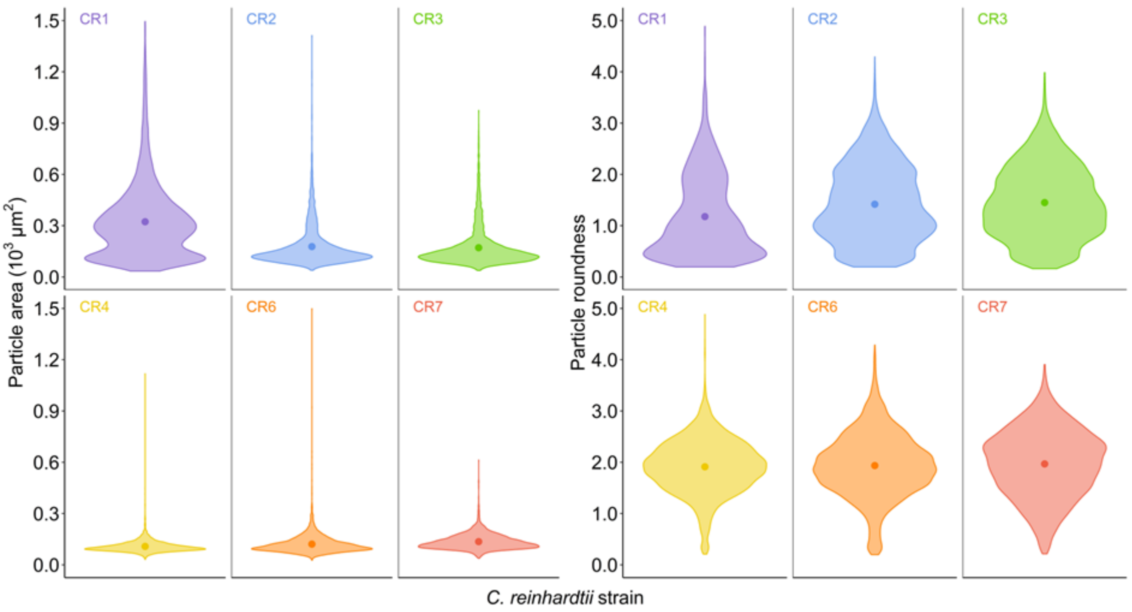


**Figure S5:** Distribution of values for 2 relevant morphological features associated to cell size (particle size) and cell shape (particle roundness) explaining most of the variance in the PCA for each *C. reinhardtii* strain (n = 5000). Mean values (dots) for the 2 morphological features were indicated within the distributions.
